# Characterization of TGFβ-induced tendon-like structure in scaffold-free three-dimensional tendon cell culture

**DOI:** 10.1101/2022.06.08.495368

**Authors:** Bon-hyeock Koo, Yeon-Ju Lee, Na Rae Park, Su-Jin Heo, David M. Hudson, Aysel A. Fernandes, Chet S. Friday, Michael W. Hast, David T. Corr, Douglas R. Keene, Sara F. Tufa, Nathaniel A. Dyment, Kyu Sang Joeng

## Abstract

Tendons transmit mechanical forces between muscle and bone. Their biomechanical function requires high tensile strength provided by highly organized collagen fibers. Tenocytes mainly drive tendon growth via extracellular matrix (ECM) production and organization. The biological mechanisms regulating tenocyte differentiation and morphological maturation have not been well-established, partly due to the lack of reliable *in vitro* systems that produce highly aligned collagenous tissues. In this study, we developed a scaffold-free, three-dimensional (3D) tendon culture system using mouse tendon cells and a differentially adherent growth channel. TGFβ treatment promoted tendon-like structure in the peripheral layer of the constructs with decreased cell density, decreased cell proliferation, increased thickness, and more elongated cells within highly aligned extracellular matrix. The constructs were used to understand the function of TGFβ signaling in tenogenic differentiation, collagen fibrillogenesis, and biomechanical properties. This scaffold-free 3D constructs system can serve as a reliable *in vitro* system to study underlying biological mechanisms that regulate cellular and matrix maturation in tendon development and growth.

## Introduction

Tendons play a critical role in the musculoskeletal system by transmitting mechanical forces between muscle and bone. Their function requires great tensile strength provided by a highly organized collagen fiber structure consisting of multiple collagen fibrils primarily composed of type I collagen (1, 2). Despite their notable tensile strength, tendons are prone to injury. Tendon injuries present a challenging clinical problem because of the slow healing process and the inability to restore original structural stability and mechanical integrity (3–6). Therefore, tendons are attractive targets for tissue engineering and regenerative medicine. However, the biological and mechanical mechanisms regulating cellular and matrix maturation remain unclear, which impedes advancing tendon tissue engineering and regenerative medicine.

Tendon maturation results in dramatic cellular and matrix changes (7, 8). Tendon fibroblasts (i.e., tenocytes) are the primary cell type that drives tendon growth via extracellular matrix (ECM) production and organization (9–11). Tenocyte differentiation is a multistep process that requires specific gene expression and unique morphological changes. Tendon progenitors are marked by the expression of *Scleraxis* (*Scx*), a basic helix-loop-helix (bHLH) transcription factor that is critical for tenocyte differentiation (12, 13). Cells at later stages of the lineage highly express type I collagen and tenomodulin (14–16). Besides molecular changes, tenocytes undergo unique morphological alterations. Younger tendons have more big and rounded cells, but fully matured tendons have more elongated tenocytes aligned longitudinally between dense and highly organized collagen fibers (8). Mature tendon cells cease proliferation and produce extracellular matrix, which results in matrix maturation and lower cell density in the postnatal tendon (7). Despite the critical role of tenocytes in tendon maturation, the biological mechanisms regulating molecular and morphological changes in tenocytes are unclear, partly due to the lack of a reliable *in vitro* system.

Standard monolayer cell cultures have been well-established and successfully used to understand the regulatory mechanisms for cellular differentiation and homeostasis in many musculoskeletal tissues such as bone, cartilage, and muscle (17–20). However, there are no established tendon cell lines. Therefore, the tendon cell culture relies on the primary cells isolated from various tendons. In addition, the tenocyte phenotype is not maintained in monolayer culture, and there is variability in cells by isolation method, mouse age, and tendon type (21, 22). It is also challenging to study ECM organization and morphological maturation of tenocytes without a 3-dimensional (3D) environment. Much of our current understanding of tendon development and postnatal tendon maturation has come from the studies of murine genetic models (23). Recent single-cell transcriptomic studies discovered tendon’s complex cellular landscape, consisting of intrinsic and extrinsic cell populations such as tendon fibroblasts, macrophages, endothelial cells, and pericytes (24–26). Tendon maturation is also regulated by multiple factors, such as biological signaling pathways and various mechanical forces (27). Due to this complexity, *in vivo* mouse models are limited in their ability to mechanistically investigate the independent biological or biomechanical factors that regulate tenocyte and matrix maturation. Therefore, developing a reliable 3D *in vitro* tendon culture system will be beneficial to overcome the limitations of the monolayer cell culture systems and animal models.

Recent scaffold-free and cell-based tissue engineering approaches have developed several unique *in vitro* tendon-like constructs. Tendon constructs have been generated from contracted or rolled-up monolayer cell culture and implanted to treat tendon injuries in small and large animal models (28–30). Another approach utilized 3D dialysis-tube-based roller culture to produce fiber-like cell aggregates presenting tendon-like organization of cells and collagen fibers (31). At the single-fiber scale, a micromold-based technique was used to generate single cellular fibers, wherein cellular self-assembly and fiber formation was directed by differentially adherent growth channels coated with fibronectin (32, 33). These scaffold-free approaches have the advantage of investigating the biological mechanism underlying tendon development, compared to scaffold-based techniques, because the construct is formed via the cells’ innate ability to self-assemble and produce extracellular matrix rather than the remodeling of a pre-existing scaffold structure. However, the application of many of these scaffold-free approaches has been limited to studies of embryonic tendon development due to high cell density and immature matrix organization.

In the current study, we developed a tissue-scale scaffold-free 3D tendon culture system by modifying the aforementioned published method for engineering single fibers via differentially-adherent growth channels (32, 33). We generated tendon-like constructs that display a tissue maturation process with key similarities to developing tendons, including decreased cell density, increased thickness, and elongated cells between highly aligned extracellular matrix. Second Harmonic Generation (SHG) microscopy confirmed the maturation of collagen fibers, and molecular analysis verified the tenogenic differentiation in the constructs. Our results suggest that the 3D tendon culture system using mouse tendon cells provides a reliable *in vitro* system to study underlying cellular and molecular mechanisms of tendon development and growth.

## Materials and Method

### Animals

All studies were approved by the Institutional Animal Care and Use Committee (IACUC) and University Laboratory Animal Resources (ULAR) at the University of Pennsylvania (Philadelphia, Pennsylvania, USA). The study was carried out in compliance with the ARRIVE guidelines (34). The mice from mixed background lines were used to generate 3D tendon constructs.

### Monolayer culture for primary tail tendon cell

Tail tendons were isolated from 25-28 days old mice. The isolated tail tendons were digested with 5ml of type I collagenase solution (2mg/mL in 1X PBS, Sigma C0130) for one hour in a 37 °C incubator by gently inverting every 10 minutes. The digested tail tendons were cultured in six-well plates with tendon growth medium (αMEM, Gibco) supplemented with 20% Fetal Bovine Serum (FBS, New Zealand origin, Sigma F8317), 1% Penicillin-Streptomycin, and 2mM L-Glutamine) for four days until 80-90% confluency. Then, tendon cells were passaged from a six-well plate to a 100mm tissue culture plate and grown in tendon growth medium for two days until 80-90% confluency was reached. Finally, the tendon cells were split into five 100mm tissue culture plates (1:5 split ratio). Cells were grown for two more days until 80-90% confluent, then harvested for 3D tendon cultures. The timeline of monolayer culture is summarized in Figure 1 (Figure 1A and 1B).

**Figure 1.**
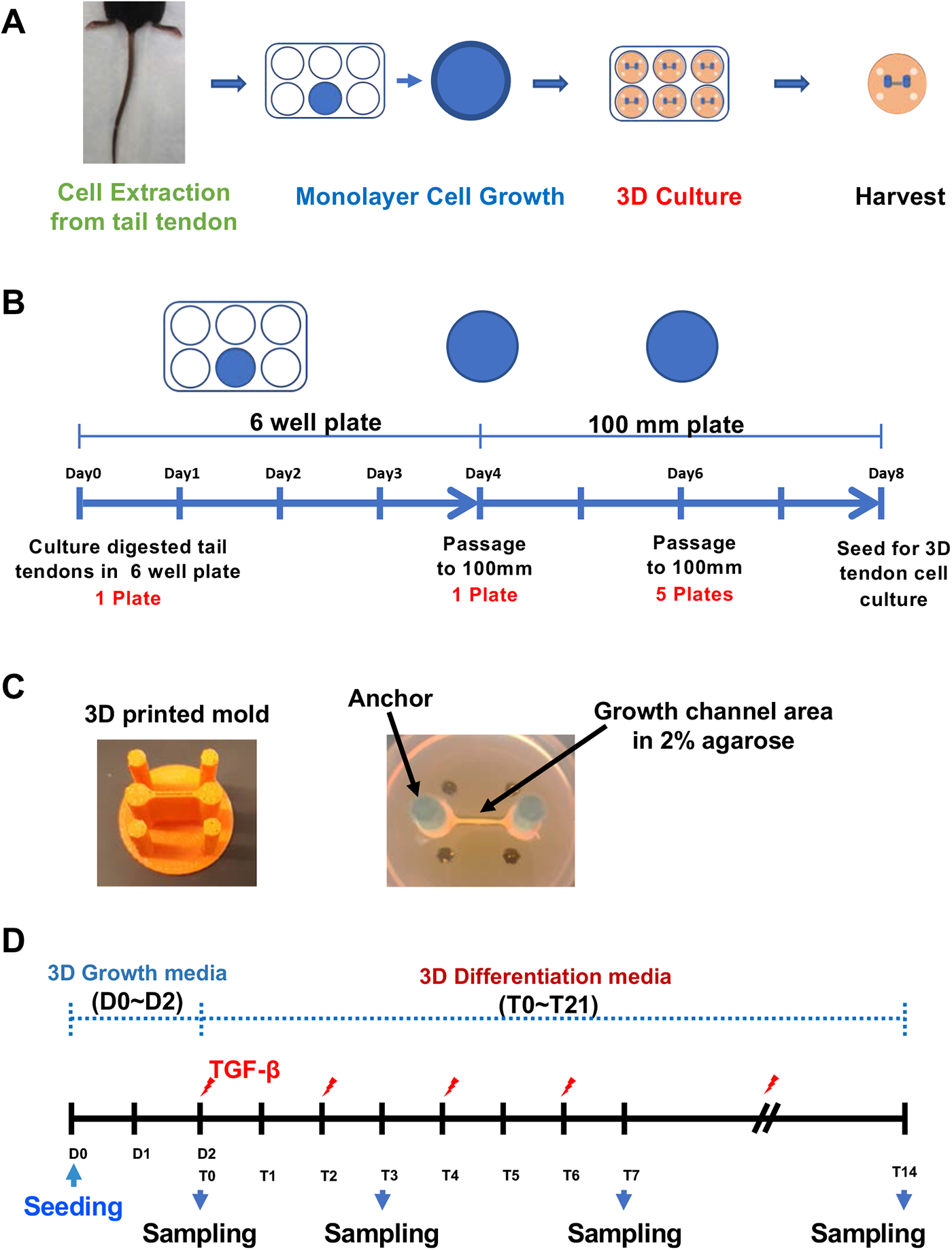
Development of 3D tendon cell culture system. (A) Three major steps of 3D tendon culture: cell extraction, cell growth in monolayer, and 3D culture. (B) The timeline of monolayer tendon cell culture after cell extraction. (C) The base of the 3D tendon cell culture consists of a growth channel area molded into 2% agarose using 3D-printed-mold and cylindrical anchors wrapped by hydrophilized PCL. Primary cells from the mouse tail tendon were seeded (2.5 x 10^6^ cells per construct) into the fibronectin-coated growth area to generate a 3D tendon construct. (D) The timeline of 3D tendon culture following monolayer culture (D, days after seeding of cells for 3D tendon culture; T, days after TGFβ treatment).

### Generation of three-dimensional (3D) tendon constructs

#### Growth channel assembly

We modified the previously published growth channel self-assembly method (32, 33). Growth channels were made using a 3D-printed mold in agarose solution (2% agarose in αMEM) on a 6-well plate (Figure 1C). The mold was removed after gelling the agarose for 30 minutes to produce a growth channel with 1.0 cm in length, 1.0 mm wide, and 3.0 mm deep. The agarose channels were sterilized by UV for 30 minutes. Anchors, 3D-printed cylinders wrapped with the hydrophilized electrospun polycaprolactone (PCL), were inserted into both ends of the growth channel area (Figure 1C), and UV sterilization was performed again for 30 minutes. The growth channels were then coated with Human plasma-derived fibronectin (0.375mg/ml in 1X PBS, Corning) and dried for 20 minutes.

#### Three-dimensional (3D) tendon cell culture

Cells were seeded (2.5 x 10∧6 cells) into the fibronectin-coated agarose growth channel area in each well. Growth media (a-MEM supplemented with 20% fetal bovine serum, 1% penicillin-streptomycin, 2mM L-Glutamine, and 50ug/ml L-Ascorbic acid) was added to the culture plate ten minutes after seeding, and the seeded culture plates were placed into a standard cell culture incubator (37°C, 5% CO2, and 95% relative humidity). Two days after seeding, growth media was replaced with differentiation media (a-MEM, supplemented with 10% fetal bovine serum, 1% penicillin-streptomycin, 2mM L-Glutamine, and 50ug/ml L-Ascorbic acid) with or without TGFβ (5ng/ml), and differentiation media was replaced every other day for the duration of time in culture. 3D tendon constructs were incubated with BrdU (5μg/ml) for two hours before harvest. The timeline of 3D tendon culture is summarized in Figure 1D.

### RT-PCR and quantitative real-time PCR (qRT-PCR)

RNA was extracted with Trizol and Direct-zol RNA miniprep kits (Zymo Research, R2050) from 3D tendon constructs. cDNA was synthesized from 500 ng of total RNA using iScript Reverse Transcription (Bio-Rad, 1708841). The qRT-PCR was performed using SYBR green master mix (Pec, 4385612) on Quantstudio 6. The sequences of primers are listed in Table 1.

**Table 1.**
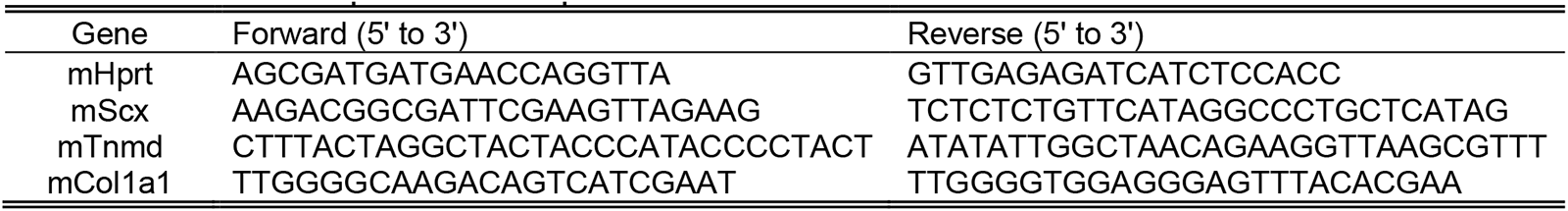
Primer sequences for qRT-PCR.

### Histological analyses

3D tendon constructs were harvested at various stages and fixed in 4% paraformaldehyde solution for one hour at 4 °C. The fixed samples were paraffin-embedded and sectioned with 6µm thickness using a microtome. These sections were used for Hematoxylin and Eosin (H&E), BrdU staining (Abcam, ab125306), DAPI staining, and TUNEL assay (Abcam, ab206386).

Nucleus shape and cell density in the outer layer of the 3D tendon structure were analyzed using DAPI-stained samples. The shape of the nucleus was characterized by the ratio of the nuclear transverse length to its length in the direction of the channel length. The fixed samples were embedded in optimal cutting temperature (OCT) compound and sectioned at 10 μm using cryostat for fluorescence imaging.

### Second Harmonic Generation

The paraffin-embedded sections were mounted with an anti-fade DAPI solution (Invitrogen, P36935) for imaging. Second Harmonic Generation (SHG) Images were generated by a Leica TCS SP8 MP multiphoton microscope (Leica Microsystems, Buffalo Grove, IL) equipped with a Coherent Chameleon Vision II Ti-Sapphire laser (Coherent, Santa Clara, CA). The images were acquired using excitation at 880 nm. DAPI images were acquired using 405 nm excitation and detection from 414-471 nm. The objective lens was a 40x/1.3 NA HC PL APO CS2 oil immersion lens. Z-stacks were acquired using a zoom of 2, a scan speed of 600 Hz, bidirectional scanning, and 6-line averaging. The lateral (x-y) pixel size was 0.0865 microns; the z-step size was 0.50 microns.

### Electron Microscopy Analysis for Collagen Fibril

The collagen fibril analysis was previously described (35). Briefly, 3D tendon constructs were harvested and fixed in the fresh buffer (1.5% glutaraldehyde/1.5% paraformaldehyde (Tousimis) with 0.05% tannic acid (Sigma) in DPBS) at 4 °C overnight. The isolated 3D tendon constructs were post-fixed in 1% OsO4 and rinsed in DMEM. The samples were then gradually dehydrated in a series of ethanol to 100%. Finally, samples were rinsed in propylene oxide, infiltrated in Spurrs epoxy, and polymerized at 70 °C overnight. A FEI G20 TEM was used for transmission electron microscopy (TEM) images which visualize transverse sections of collagen fibrils at multiple magnifications. Collagen fibril diameter was measured using ImageJ software. TEM images of six constructs per group were analyzed. Total of 3452 fibrils in vehicle-treated constructs and 3507 fibrils in TGFβ-treated constructs were counted.

#### Pyridinoline cross-linking analysis

Multiple tendon constructs (n = 2-6) from each time point were pooled for cross-linking analysis. The pyridinoline cross-link content was determined for each pooled group of tendons by HPLC after acid hydrolysis in 6 N HCl for 24 hrs at 108°C. Dried samples were dissolved in 1% (v/v) n-heptafluorobutyric acid, and hydroxylysine pyridinoline (HP) content was quantified by fluorescence monitoring during reverse-phase HPLC as previously described (36).

### Uniaxial Biomechanical testing

Uniaxial testing was performed similarly to previously established protocols (37–39). All specimens were kept frozen at -20°C until the day of testing. After thawing, a custom laser device was used to measure the cross-sectional area (mm^2^). The ends of each specimen were placed between two sandpaper tabs and held together with cyanoacrylate glue to prevent specimen slippage with the grips during the mechanical test. Samples were gripped with custom aluminum fixtures and mounted onto a universal test frame (Instron 5542, Instron Inc., Norwood, MA) with a 10N load cell. All testing was conducted in a phosphate-buffered saline bath maintained at room temperature (23°C). For biomechanical evaluation, each sample was preloaded to 0.02 N, followed by ten cycles of preconditioning between 0.02 and 0.04 N. The specimens then underwent a quasi-static ramp to failure at a displacement rate of 0.01 mm/s. Force, displacement, and time data were collected at 100Hz. Force-displacement curves were analyzed to quantify tissue stiffness (N/mm, defined as the slope of the linear region) and ultimate force. Force and displacement data were normalized by specimen CSA and gauge length, respectively, to convert to stress and strain. The resulting stress-strain curves were assessed to determine elastic modulus (MPa, defined as the slope of the linear region) and failure stress (MPa). Work to failure (mJ) and toughness (mJ/m^3^) were determined by calculating the areas under the force-displacement and stress-strain curves, respectively.

### Statistical analysis

Results are expressed as mean ± SD. At least three 3D tendon constructs per group were analyzed. Differences between values were analyzed by Student’s t-test. P<0.05 is considered significant.

## Results

### TGFβ treatment induced tendon-like tissue maturation in 3D tendon constructs

The self-assembly of seeded cells in the fibronectin-coated channel formed the tendon-like constructs within 24 hours after the seeding. However, the thickness of the constructs was not maintained with regular growth media (Figure 2A, vehicle). Microscopic analysis at higher magnification showed many cells budding out from the surface of the vehicle-treated constructs at T4 and T7 (Figure 2B, upper panels). To analyze the structure of the 3D tendon constructs, we collected the constructs at different time points and performed histological analysis using H&E-stained longitudinal sections (Figure 2C). Lower magnification images confirmed that the thickness of the vehicle-treated construct was significantly reduced at T14 (Figure 2C, 20X whole, left panels). We also found that the 3D tendon constructs contained two cell layers, inner (Figure 2C, yellow asterisks) and peripheral (Figure 2C, red arrows) layers. Dramatic decrease in inner layer thickness was observed in vehicle-treated constructs at T14 (Figure 2C, 20X whole, left panels). The quantification results verified the gradual decrease in the inner layer thickness of vehicle-treated constructs (Figure 2D, upper left panel, white bar). There was no significant change in cell density of the inner layer in vehicle-treated constructs (Figure 2D, upper right panel, white bar). Lower magnification images showed a slight reduction in peripheral layer thickness of vehicle-treated constructs (Figure 2C, 20X whole, vehicle), which is confirmed by the higher magnification image (Figure 2C, 40X peripheral, vehicle). In our quantification analysis, we observed slightly reduced peripheral layer thickness in vehicle-treated constructs (Figure 2D, bottom left panel, white bars). A gradual reduction in cell density of the inner layer was also observed in vehicle-treated constructs ((Figure 2D, bottom right panel, white bars). Previous studies suggested that additional growth factors could be essential for maintaining the engineered tendon constructs (29, 31, 32). Transforming growth factor-beta (TGFβ) is a well-known growth factor and critical for tendon development, maturation, healing, and tenogenic differentiation *in vivo* and *in vitro* (16, 31, 40–47). These prior findings prompted us to use TGFβ in our 3D tendon culture. Interestingly, the thickness of the construct was relatively well-preserved with TGFβ treatment compared to that of the vehicle-treated construct at T14 (Figure 2A). Microscopic analysis at higher magnification further confirmed that TGFβ treatment preserved the thickness of constructs and maintained the smooth surface without the budding cells at earlier stages, such as T4 and T7 (Figure 2B, bottom panels). Histological analysis verified that the thickness of the TGFβ-treated construct was relatively preserved when compared to vehicle-treated constructs (Figure 2C, 20X whole, right panels). The inner layer thickness of TGFβ-treated constructs was also gradually reduced (Figure 2C, 20X whole) at T14, which is confirmed by quantification (Figure 2D, upper left panel, gray bars). However, the inner layer of TGFβ-treated construct was significantly thicker than that of vehicle-treated constructs at T14. TGFβ-treated constructs displayed higher cell density in the inner layer at T3 and T7 when compared to vehicle-treated constructs, but there was no difference at T14 (Figure 2D, upper right panel). The peripheral layer of TGFβ-treated constructs became much thicker at T14 when compared to early stages such as T3 and T7 (Figure 2C, 20X whole and 40X peripheral, TGFβ+). The quantification results revealed dramatically increased peripheral layer thickness of TGFβ-treated constructs at T7 and T14 (Figure 2D, bottom left panel, gray bars). Interestingly, the higher magnification image shows that the peripheral layer of the TGFβ-treated constructs presented a tendon-like maturation, including elongated cells between a highly aligned extracellular matrix, increased thickness, and decreased cell density (Figure 2C, 40X peripheral, TGFβ+). Our quantification results verified the dramatic reduction in the peripheral layer cell density of TGFβ-treated constructs throughout the stages (Figure 2D, bottom right panel, gray bars). Overall, these findings strongly suggest that TGFβ is indispensable for promoting proper aggregation and self-assembly of the 3D tendon constructs, and TGFβ treatment induces tendon-like tissue maturation in the peripheral layer of the constructs.

**Figure 2.**
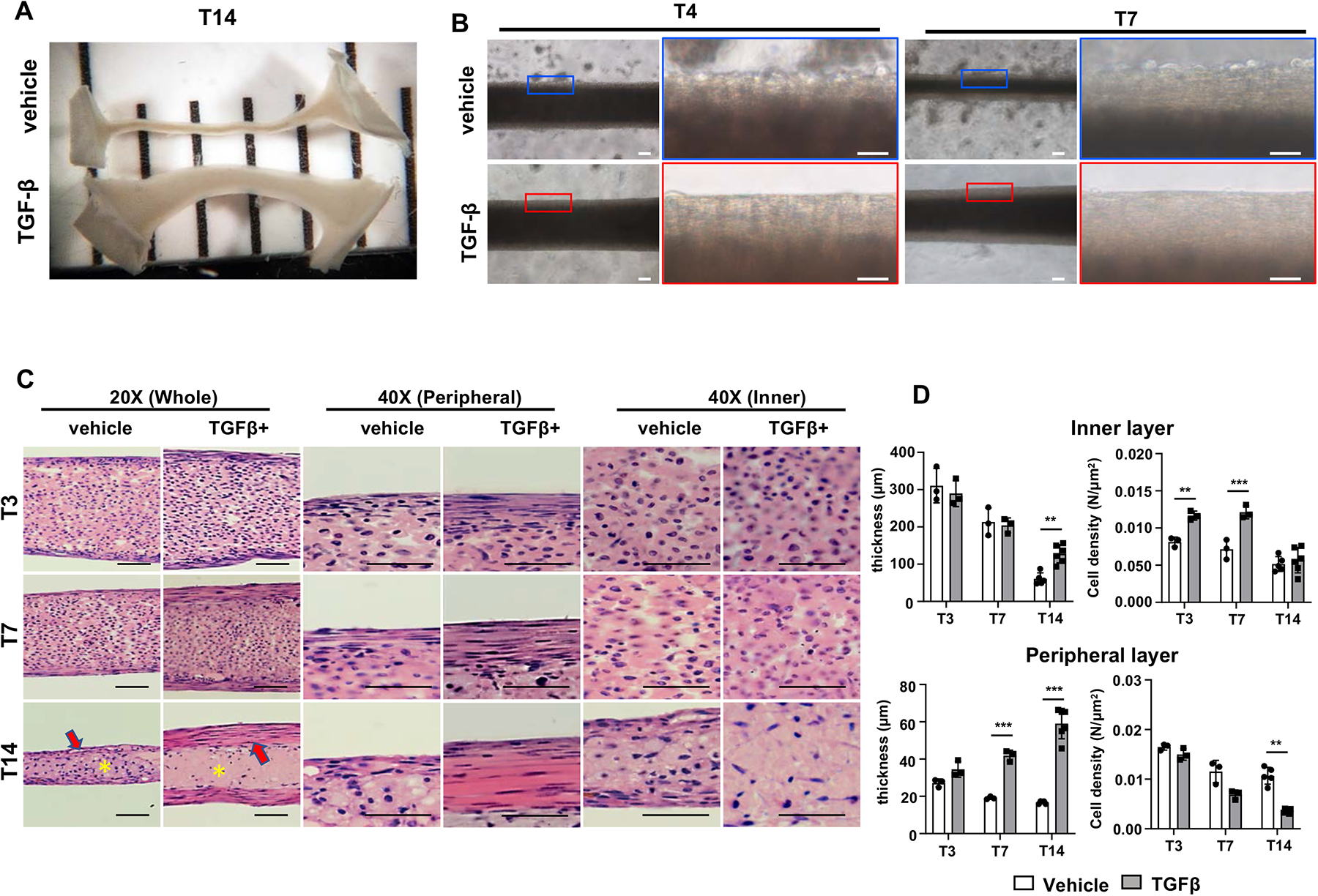
Histological analyses of the 3D tendon constructs. (A) Dissecting microscope images of 3D tendon constructs at 14 days after TGFβ or vehicle treatment (T14). (B) Higher magnification images of 3D tendon constructs at T4 and T7 after TGFβ or vehicle treatment. Scale bars in lower magnification images indicate 100μm, and scale bars in higher magnification images indicate 50μm. (C) H&E-stained longitudinal sections from vehicle- and TGFβ-treated 3D tendon constructs at each stage (T3, T7, and T14). Scale bars in lower magnification images indicate 100μm, and scale bars in higher magnification images indicate 20μm. Yellow asterisks indicate inner layers, and red arrows indicate peripheral layers (D) Quantification results of thickness and cell density of inner and peripheral layer (** indicates P<0.01 and *** indicates P<0.001, n=3∼5 constructs per group).

### TGFβ treatment increased cell proliferation in 3D tendon constructs

The decreased cell density in the peripheral layer of 3D tendon constructs prompted us to investigate cellular proliferation and apoptosis. To study cellular proliferation, we performed BrdU staining with longitudinal sections of 3D tendon constructs (Figure 3A). The quantification results show that both vehicle-and TGFβ-treated constructs exhibit gradual reduction in cellular proliferation in both layers with time in culture (Figure 3B). However, TGFβ-treated constructs display significantly higher proliferation rates at T7 and T14. To examine apoptosis, we performed the Tunnel assay using longitudinal sections of 3D tendon constructs (Figure 3C). In the peripheral layer, vehicle-treated constructs display a higher apoptosis rate than TGFβ-treated constructs at T3, but no difference was observed at T7 and T14 (Figure 3D, left panel). In the inner layer, both constructs present high apoptosis rates at T3 and T7, but the apoptosis is significantly reduced at T14 (Figure 3D, right panel). Interestingly, TGFβ-treated constructs show a significantly higher apoptosis rate at T14 when compared to vehicle-treated constructs. Overall, TGFβ treatment increased cellular proliferation in both peripheral and inner layers but only prevented cellular apoptosis in the peripheral layer at early stages.

**Figure 3.**
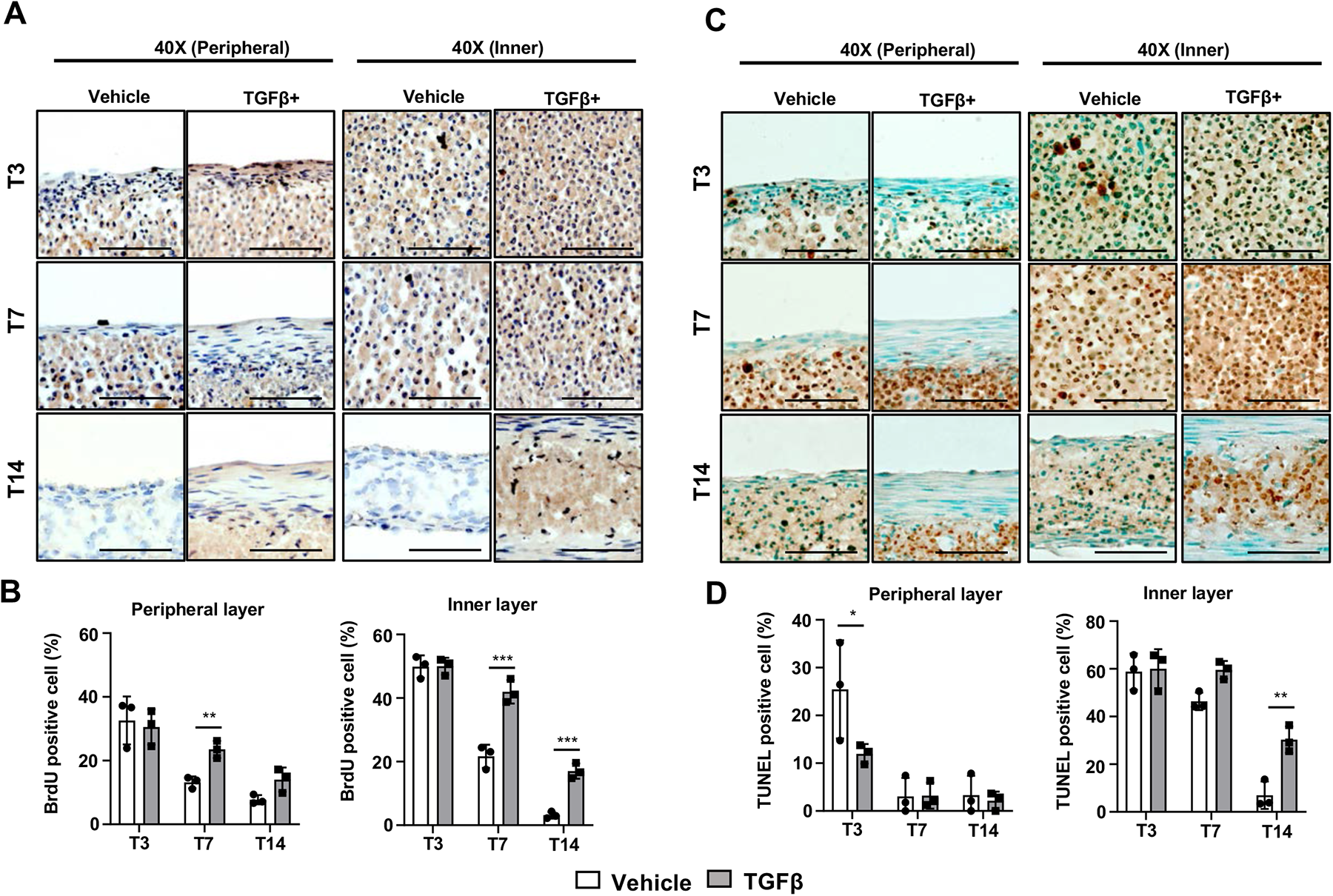
Cellular proliferation and apoptosis in the 3D tendon constructs. (A) BrdU staining of peripheral and inner layers in vehicle- and TGFβ-treated 3D tendon constructs at each stage (T3, T7, and T14). Scale bars indicate 20μm. (B) Quantification results of BrdU-positive cells in peripheral and inner layers of 3D tendon constructs (** indicates P<0.01 and *** indicates P<0.001, n=3). (C) TUNEL assay of peripheral and inner layers in vehicle- and TGFβ-treated 3D tendon constructs at each stage (T3, T7, and T14). (D) Quantification results of Tunnel-positive cells in peripheral and inner layers of 3D tendon constructs (* indicates P<0.05 and ** indicates P<0.01, n=3 constructs per group).

### TGFβ increased collagen fiber formation but reduced collagen fibril diameter

Collagen fiber formation is one of the major features of tendon maturation. Tendon-like tissue maturation of the peripheral layer of TGFβ-treated constructs prompted us to assess collagen fiber formation using Second Harmonic Generation (SHG) microscopy with longitudinal sections of the 3D tendon constructs (Figure 4A). A gradual increase in SHG signals was observed in peripheral layers of both vehicle and TGFβ-treated constructs, but TGFβ-treated constructs contained more fibers with stronger SHG signal. Specifically, no SHG signal was detected in the peripheral layer of vehicle-treated constructs at T3, and only a weak signal was detected at T7 and T14. SHG signal was also undetectable in the TGFβ-treated constructs at T3, but strong linear SHG signals were detected at T7 and T14, suggesting the TGFβ treatment caused the production of mature, well-organized collagen matrix at later stages. No SHG signals were detected in the inner layers of both constructs.

**Figure 4.**
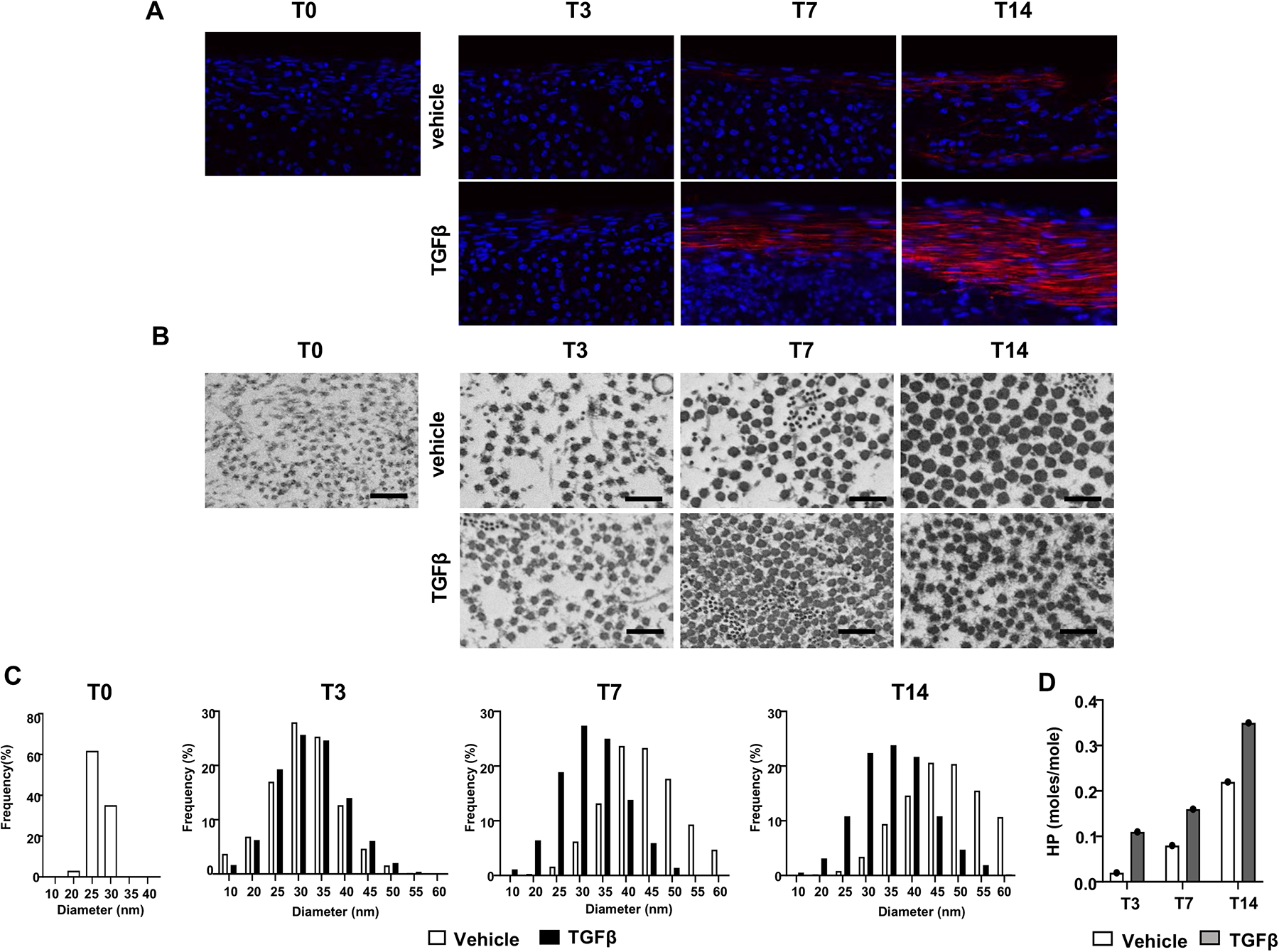
Collagen fiber formation, modification and fibrillogenesis in the 3D tendon constructs. (A) Second Harmonic Generation (SHG) microscopy images (collagen fiber formation) of peripheral layers of vehicle-and TGFβ-treated 3D tendon constructs at each stage (T0, T3, T7, and T14). (B) Transmission electron microscopy (TEM) microscopy images of peripheral layers of vehicle- and TGFβ-treated 3D tendon constructs at each stage (T0, T3, T7, and T14). (C) Quantification of collagen fibril diameter using TEM images from vehicle- and TGFβ-treated 3D tendon constructs at each stage (n=3 constructs per group at each stage). (D) hydroxylysylpyridinoline level of vehicle- and TGFβ-treated 3D tendon constructs at each stage (combined samples (2-4 constructs per group) were used for the analysis).

We further examined collagen fibril ultrastructure and cross-linking because collagen fibrillogenesis and modification are critical for tendon maturation. Transmission electron microscopy (TEM) was performed using 3D tendon constructs to measure collagen fibril diameter. Transmission electron microscopy (TEM) images visualize transverse sections of collagen fibrils, and both constructs showed gradual growth of collagen fibrils diameter throughout the stages (Figure 4B). The quantification results demonstrated the gradual growth of collagen fibril in vehicle-treated constructs (Figure 4C, white bars). Specifically, at T0, the constructs displayed a narrow range of collagen fibril sizes, with the greatest proportion of collagen fibrils at a diameter of 25 nm. The size (diameter) of the greatest proportion of fibrils gradually increased to 30 at T3, 40 at T7, and 45 at T14. TGFβ-treated constructs showed a similar growth of collagen fibril diameter at T3 when compared to vehicle-treated constructs (Figure 4C, black bar), but the growth in fibril diameter was surprisingly inhibited by TGFβ treatment, indicated by the greatest proportion of collagen fibril at a diameter of 30 for T7 and 35 for T14. Collagen cross-linking analyses were performed using 3D tendon constructs to assess the status of collagen modification. The cross-linking analysis revealed an increase in hydroxylysylpyridinoline in TGFβ-treated tendon constructs at all stages (Figure 4D). It should be noted that while hydroxylysylpyridinoline provides a useful measure of stable (irreversible) cross-links, it is not a measure of the total cross-links. Overall, our collagen analyses indicate that the peripheral layer of the 3D tendon construct undergoes tendon-like structural maturation, and TGFβ enhances collagen fiber formation and cross-linking, but inhibits the lateral growth of collagen fibril.

### TGFβ induced morphological maturation and tenogenic differentiation

To investigate cellular phenotypes in 3D tendon constructs, we first examined the morphological changes of cells in the 3D tendon construct (Figures 5A and 5B). We measured the ratio of the nuclear transverse length to its length in the direction of the channel length. In this measurement, a value of 1.0 would indicate a perfect circle, and 0 would be a line in the direction of the channel length. Therefore, nuclear height/width ratios closer to 1.0 indicates a rounder cell shape, whereas values less than 1 indicate a cell shape more elongated in the direction of construct length, with smaller values indicating greater cellular elongation. Gradual cell elongation was observed in the peripheral layer of both vehicle- and TGFβ-treated constructs, with the TGFβ-treated constructs exhibiting more elongated cells than vehicle-treated constructs throughout the stages (Figure 5A). Specifically, the cells in the 3D tendon constructs were relatively oval-shaped at T0, which is evident with the major distributed values between 0.3 and 0.5 (Figure 5A, T0 peripheral layer). Vehicle-treated constructs exhibited minimal morphological changes at T3, and the cells became elongated at T7 and T14, which is evident with the major distributed values between 0.2 and 0.3 (Figure 5A, white bar). The cells in TGFβ-treated constructs presented dramatic morphological changes at T3 and T7, which is evident with most of the values distributed between 0.1 and 0.3 (Figure 5A, black bar). The cells became further elongated at T14, with the greatest proportion of cells at a ratio of 0.1. Within the inner layer, the cells only show morphological changes between T0 and T3, but no further changes were observed. Additionally, TGFβ treatment does not appear to affect the morphological maturation of the inner layer cells (Figure 5B). These data suggest that the cells in the peripheral layer undergo morphological maturation, and TGFβ can be used to enhance this morphological maturation.

**Figure 5.**
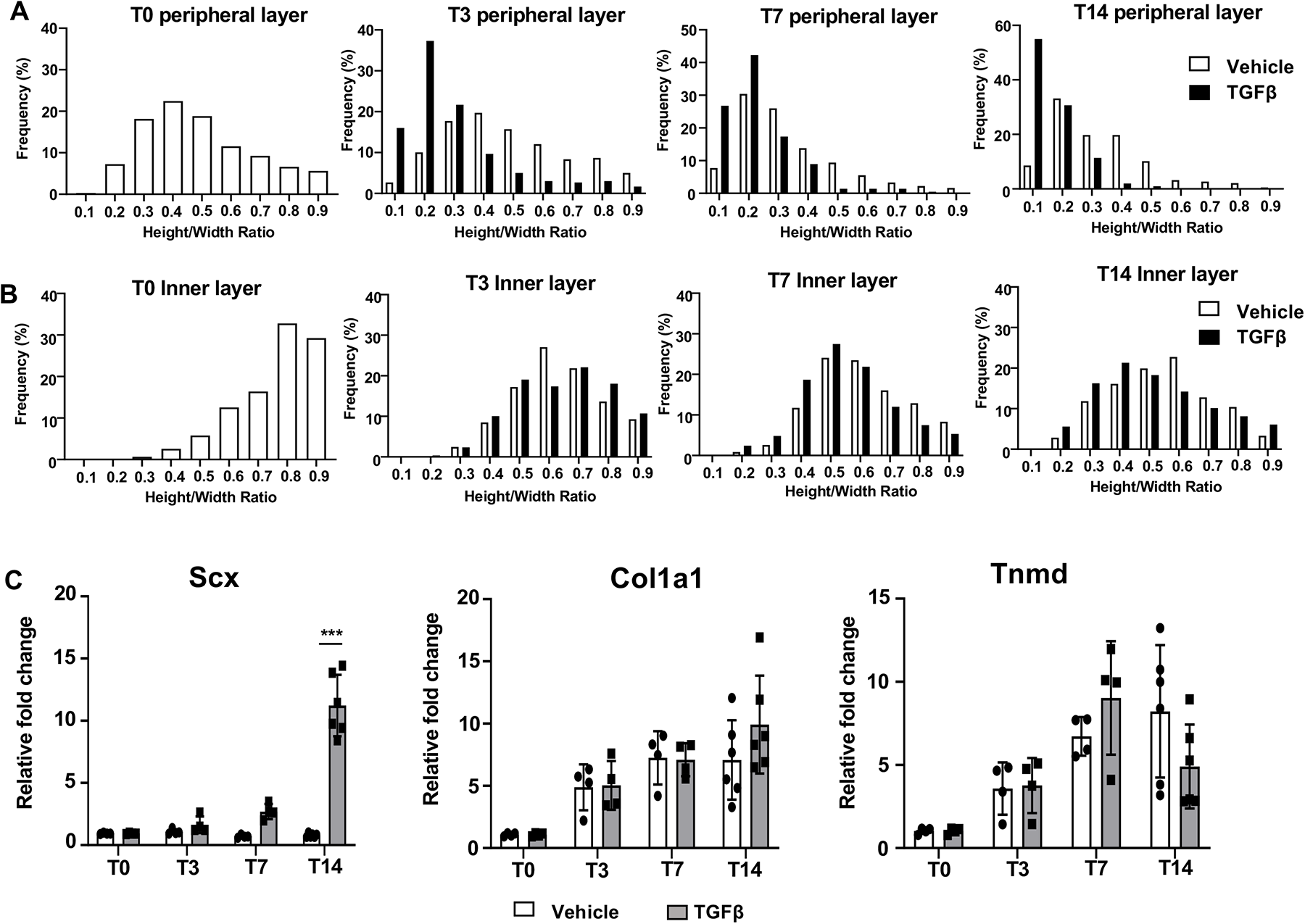
Morphological and molecular changes of cells in the 3D tendon constructs. The ratio of the nuclear transverse length to its length in the direction of the channel length of cells in peripheral (A) and inner (B) layers (values closer to 1.0 indicates a rounder cell shape, and values closer to 0 indicate a more elongated cell shape). n=3 constructs per group. (C) Quantitative real-time PCR (qRT-PCR) results for tenogenic gene expression in 3D tendon constructs, including Scleraxis (Scx), Tenomodulin (Tnmd), and Collagen type I (Col1). *** indicates P<0.001 (n=4-6 constructs per group).

To assess tenogenic differentiation, we examined the expression of three different tenogenic gene markers, *Scleraxis (Scx)*, *Tenomodulin (Tnmd)*, and Collagen type 1 (*Col1a1*) (Figure 5C). First, TGFβ treatment significantly induced the expression of *Scleraxis*, a critical transcription factor for tenogenic differentiation (Figure 5C, Scx). TGFβ treatment increased the expression of *Scleraxis* at T14 by 10-fold compared to T0, while *Scleraxis* expression was not changed at all stages of vehicle-treated constructs. Collagen type 1 and *Tenomodulin*, relatively later tenogenic markers, were significantly increased in later stages in both vehicle- and TGFβ-treated constructs, but no significant changes were observed between the two treatments (Figure 5C, Col1a1 and Tnmd). These data suggest that TGFβ may promote tenogenic cell differentiation in the 3D tendon constructs.

### TGFβ treatment enhanced the mechanical properties of 3D tendon constructs

We performed uniaxial biomechanical testing with samples isolated at T14. Vehicle-treated tendon constructs were very thin and extremely delicate, so they did not possess sufficient mechanical integrity to allow for biomechanical characterization. Indeed, all vehicle-treated samples broke prior to the completion of mechanical testing. On the other hand, TGFβ-treated samples were thicker and stronger, so mechanical testing was possible. The cross-sectional area of TGFβ-treated constructs were similar to that of the Achilles tendons from one-month-old mice (35, 48) and exhibited moduli similar to those of Achilles tendons from neonatal mice at day 4 and day 7 (48). The measured mechanical property values are summarized in Table 2.

**Table 2.** Mechanical properties of TGFβ−treated 3D tendon constructs at T14 (n=7, value ± standard deviation)

|  |  |
| --- | --- |
| Cross-Sectional Area (mm<sup>2</sup>) | 0.17 ± 0.10 |
| Max Load(N) | 0.43 ± 0.14 |
| Stiffness(N/mm) | 0.23 ± 0.07 |
| Max Stress (MPa) | 3.52 ± 2.43 |
| Modulus (MPa) | 20.55 ± 9.92 |
| Energy To Failure (mJ) | 0.50 ± 0.26 |
| Max Strain (mm/mm) | 0.19 ± 0.06 |
| Toughness (mJ/m<sup>3</sup>) | 0.39 ± 0.38 |

## Discussion

In this study, we successfully generated *in vitro* 3D constructs using mouse tail tendon cells. TGFβ treatment induced tendon-like structure in the peripheral layer of constructs, exhibiting decreased cell density and cell proliferation, increased thickness, and elongated cells between highly aligned extracellular matrix. We also characterized collagen fibrillogenesis, tenogenic differentiation, and biomechanical properties of the 3D tendon constructs. These well-characterized constructs will be a valuable tool to investigate cellular and molecular mechanisms regulating tendon development and maturation *in vitro* and will also contribute to the understanding of scaffold-free bioengineered graft models.

Previous studies suggested the potential requirement of growth factors for the formation of engineered tendon constructs (29, 31, 32). Our results suggest that TGFβ is essential for successful 3D tendon culture. TGFβ treatment maintains the thickness of the 3D tendon due to increased inner and peripheral layer thickness compared to vehicle-treated control constructs. The decreased cell density with increased thickness in the peripheral layer suggests that TGFβ treatment induces the production of extracellular matrix. TGFβ dramatically induced the expression of *Scleraxis*, the critical transcription factor for tenogenic differentiation. However, the expression of *Col1a1* and *Tenomodulin* were not changed by TGFβ treatment. These data suggest that cells undergo early tenogenic differentiation without further molecular maturation with TGFβ treatment. We carefully speculate that this early arrested cell fate could explain the smaller collagen fibril diameters in TGFβ-treated tendon constructs when compared to control constructs. Besides TGFβ, the functions of several growth factors, such as BMPs, CTGF, and FGFs, have been investigated in physiological and pathological tendon conditions using *in vivo* and *in vitro* models (16, 49–52). Upon the existence of recombinant proteins of these growth factors, our 3D tendon constructs can be used to test the function of these growth factors in tenogenic cell differentiation and matrix maturation.

The success of the initial monolayer culture is critical for generating 3D tendon constructs. In our preliminary study, the tail tendon cells from mice older than one month grew too slowly in monolayer culture, so we could not obtain enough cells for 3D tendon culture in the proper timeline. However, tail tendons from younger mice, e.g., 25-28 days old, were able to produce sufficient cell numbers for the studies herein. We assume that this could be attributed to tendons from young mice having more progenitor cells which can proliferate in *in vitro* monolayer cell culture. Interestingly, the origin of FBS was also critical for the monolayer tendon cell cultures. Our preliminary studies revealed that the primary tendon cells in the monolayer culture with US-originated FBS grew too slowly, so these cells could not be used for 3D tendon culture. However, the cells with New Zealand-originated and Australian-originated FBS grew fast enough to follow the cell culture timeline. Therefore, we only isolated tail tendon cells from younger than one-month-old mice and used New Zealand- or Australian-originated FBS for the current study.

We only used mouse tail tendon cells for the current study because we could easily isolate enough cells for 3D tendon culture. We can generate six 3D tendon constructs from one mouse tail by strictly following the timelines of the monolayer cell culture demonstrated in the Method section. In our experience, neither the Achilles nor Patellar tendons from one mouse could provide enough cells, even at the monolayer cell culture stage. However, we believe that combining multiple tendons from several mice would provide sufficient cells for the 3D tendon culture using Achilles or Patellar tendons. One of the future research directions using this platform is to compare the 3D tendon constructs generated using cells from different tendons (e.g., flexor, extensor) to identify whether the tissue origin of cells influences the development, structure, and function of engineered 3D tendon constructs (53).

In our 3D tendon culture systems, the peripheral layer of the constructs showed a tendon-like structure, but the central regions of the construct exhibited degeneration, and eventually, the inner core tissues reduced their thickness in both vehicle- and TGFβ-treated constructs (Figure 2C). We speculate that this is due to the nutritional deficiency in the center region. This phenomenon was also described in the previous culture study (31). The potential solution for core cell degeneration can be a combined treatment of biomechanical force and growth factors. Indeed, TGFβ treatment prevented the reduction of inner core thickness when compared to the vehicle treatment, but the inner layer still does not show a tendon-like structure. Additionally, our prior scaffold-free work at the single-fiber scale, which is not subject to such diffusion-based nutritional deficiency, has demonstrated that cyclic tensile loading can be utilized to improve the tensile strength as well as elastic modulus and to increase nuclear elongation of engineered tendon fibers (33). Hence, we are currently developing a bioreactor in which we can apply cyclic loading to the 3D tendon constructs. Investigating the combinatory effects of the cyclic loading and growth factors on the maturation of 3D tendon constructs will be an interesting future direction.

Although the cross-sectional area of the tendon constructs is similar to the mouse Achilles tendons at four weeks of age, overall mechanical properties are more similar to the neonatal mouse model (35, 48). This mechanical test data is consistent with our collagen fibril diameter analysis showing the diameter range is between 0nm to 60nm, which is similar to the Achilles tendons at four days of age (48). However, our constructs contain two layers of tissue structure, so the measured mechanical properties do not fully represent the properties of peripheral tendon-like structures. Therefore, fully matured uni-structured tendon constructs need to be generated to understand the precise mechanical properties of the constructs.

Our scaffold-free 3D tendon culture can be a reliable *in vitro* system to investigate the underlying cellular and molecular mechanisms regulating tendon development and maturation. The future direction of this study will be inducing the cells into a further matured stage expressing higher levels of Col1a1 and Tenomodulin with increased collagen fibril diameters. Further maturation of tendon constructs could improve the mechanical properties of current 3D tendon constructs. In addition, genetic manipulation is feasible for our 3D tendon culture using an adenovirus system, which suggests the potential usage of our constructs for *in vitro* genetic studies for specific gene functions in tendon maturation. We can also utilize our constructs for the screening of tenogenic factors using small molecule library. Application of our constructs for *in vitro* studies can provide unique insight and understanding of biological mechanisms in tendon and bioengineered graft models.

## Abbreviations

3D: three dimensional
Scx: Scleraxis
Col1: Collagen type I
Tnmd: Tenomodulin
DAPI: 4′,6-diamidino-2-phenylindole
FBS: Fetal Bovine Serum
ECM: Extracellular matrix
SHG: Second Harmonic Generation (SHG)
PBS: Phosphate-buffered saline
PCL: Polycaprolactone

## Data Availability Statement

The data that support the findings of this study are available in the methods of this article.

## Author Contributions

Study conception and design: KSJ, NAD, DTC, YJL, SJH; Acquisition of data: BK, YJL, NRP, MWH, CSF, DMH, DRK, AAF, SFT; Analysis and interpretation of data: BK, YJL, KSJ, DTC, NAD; Drafting of the manuscript: BK, YJL, DTC, KSJ.

## Acknowledgement

We would like to thank Penn Center for Musculoskeletal Disorders (PCMD) Histology and Biomechanics Core for technical assistance with the histology and 3D printing, respectively. We also thank Cell and Developmental Biology (CDB) Microscopy Core for technical assistance with confocal microscopy. This work was supported in part by a grant from the U.S. National Institutes of Health (NIH), National Institute of Arthritis and Musculoskeletal and Skin Diseases (KSJ: R01 AR079486). The PCMD Cores were supported by a grant from NIH/NIAMS P30AR069619.

## References

1. Wang, J. H.-C., Guo, Q., and Li, B. (2012) Tendon biomechanics and mechanobiology--a minireview of basic concepts and recent advancements. J. hand Ther. Off. J. Am. Soc. Hand Ther. 25, 133–140; quiz 141

2. Zhang, G., Young, B. B., Ezura, Y., Favata, M., Soslowsky, L. J., Chakravarti, S., and Birk, D. E. (2005) Development of tendon structure and function: regulation of collagen fibrillogenesis. J. Musculoskelet. Neuronal Interact. 5, 5–21

3. Galatz, L. M., Gerstenfeld, L., Heber-Katz, E., and Rodeo, S. A. (2015) Tendon regeneration and scar formation: The concept of scarless healing. J. Orthop. Res. 33, 823–831

4. Andarawis-Puri, N., Flatow, E. L., and Soslowsky, L. J. (2015) Tendon basic science: Development, repair, regeneration, and healing. J. Orthop. Res. 33, 780–784

5. Nichols, A. E. C., Best, K. T., and Loiselle, A. E. (2019) The cellular basis of fibrotic tendon healing: challenges and opportunities. Transl. Res. 209, 156–168

6. Bey, M. J. and Derwin, K. A. (2012) Measurement of in vivo tendon function. J. shoulder Elb. Surg. 21, 149–157

7. Grinstein, M., Dingwall, H. L., O’Connor, L. D., Zou, K., Capellini, T. D., and Galloway, J. L. (2019) A distinct transition from cell growth to physiological homeostasis in the tendon. Elife 8

8. Chen, J., Zhang, W., Liu, Z., Zhu, T., Shen, W., Ran, J., Tang, Q., Gong, X., Backman, L. J., Chen, X., Chen, X., Wen, F., and Ouyang, H. (2016) Characterization and comparison of post-natal rat Achilles tendon-derived stem cells at different development stages. Sci. Rep. 6, 22946

9. Dex, S., Lin, D., Shukunami, C., and Docheva, D. (2016) Tenogenic modulating insider factor: Systematic assessment on the functions of tenomodulin gene. Gene 587, 1–17

10. Sharma, P. and Maffulli, N. (2006) Biology of tendon injury: healing, modeling and remodeling. J. Musculoskelet. Neuronal Interact. 6, 181–190

11. Nichols, A. E. C., Best, K. T., and Loiselle, A. E. (2019) The cellular basis of fibrotic tendon healing: challenges and opportunities. Transl. Res. 209, 156–168

12. Murchison, N. D., Price, B. A., Conner, D. A., Keene, D. R., Olson, E. N., Tabin, C. J., and Schweitzer, R. (2007) Regulation of tendon differentiation by scleraxis distinguishes force-transmitting tendons from muscle-anchoring tendons. Development 134, 2697– 2708

13. Schweitzer, R., Chyung, J. H., Murtaugh, L. C., Brent, A. E., Rosen, V., Olson, E. N., Lassar, A., and Tabin, C. J. (2001) Analysis of the tendon cell fate using Scleraxis, a specific marker for tendons and ligaments. Development 128, 3855–3866

14. Docheva, D., Hunziker, E. B., Fässler, R., and Brandau, O. (2005) Tenomodulin is necessary for tenocyte proliferation and tendon maturation. Mol. Cell. Biol. 25, 699–705

15. Léjard, V., Brideau, G., Blais, F., Salingcarnboriboon, R., Wagner, G., Roehrl, M. H. A., Noda, M., Duprez, D., Houillier, P., and Rossert, J. (2007) Scleraxis and NFATc regulate the expression of the pro-alpha1(I) collagen gene in tendon fibroblasts. J. Biol. Chem. 282, 17665–17675

16. Huang, A. H., Lu, H. H., and Schweitzer, R. (2015) Molecular regulation of tendon cell fate during development. J. Orthop. Res. Off. Publ. Orthop. Res. Soc. 33, 800–812

17. Gordon, J. A. R., Tye, C. E., Sampaio, A. V, Underhill, T. M., Hunter, G. K., and Goldberg, H. A. (2007) Bone sialoprotein expression enhances osteoblast differentiation and matrix mineralization in vitro. Bone 41, 462–473

18. Sudo, H., Kodama, H. A., Amagai, Y., Yamamoto, S., and Kasai, S. (1983) In vitro differentiation and calcification in a new clonal osteogenic cell line derived from newborn mouse calvaria. J. Cell Biol. 96, 191–198

19. Langley, B., Thomas, M., Bishop, A., Sharma, M., Gilmour, S., and Kambadur, R. (2002) Myostatin inhibits myoblast differentiation by down-regulating MyoD expression. J. Biol. Chem. 277, 49831–49840

20. Buxton, A. N., Bahney, C. S., Yoo, J. U., and Johnstone, B. (2011) Temporal exposure to chondrogenic factors modulates human mesenchymal stem cell chondrogenesis in hydrogels. Tissue Eng. Part A 17, 371–380

21. Yao, L., Bestwick, C. S., Bestwick, L. A., Maffulli, N., and Aspden, R. M. (2006) Phenotypic drift in human tenocyte culture. Tissue Eng. 12, 1843–1849

22. Nichols, A. E. C., Muscat, S. N., Miller, S. E., Green, L. J., Richards, M. S., and Loiselle, A. E. (2021) Impact of isolation method on cellular activation and presence of specific tendon cell subpopulations during in vitro culture. FASEB J. Off. Publ. Fed. Am. Soc. Exp. Biol. 35, e21733

23. Mienaltowski, M. J. and Birk, D. E. (2014) Mouse models in tendon and ligament research. Adv. Exp. Med. Biol. 802, 201–230

24. Kendal, A. R., Layton, T., Al-Mossawi, H., Appleton, L., Dakin, S., Brown, R., Loizou, C., Rogers, M., Sharp, R., and Carr, A. (2020) Multi-omic single cell analysis resolves novel stromal cell populations in healthy and diseased human tendon. Sci. Rep. 10, 13939

25. Yin, Z., Hu, J., Yang, L., Zheng, Z.-F., An, C., Wu, B., Zhang, C., Shen, W.-L., Liu, H., Chen, J., Heng, B. C., Guo, G., Chen, X., and Ouyang, H.-W. (2016) Single-cell analysis reveals a nestin^+^ tendon stem/progenitor cell population with strong tenogenic potentiality. Sci. Adv. 2, e1600874

26. De Micheli, A. J., Swanson, J. B., Disser, N. P., Martinez, L. M., Walker, N. R., Oliver, D. J., Cosgrove, B. D., and Mendias, C. L. (2020) Single-cell transcriptomic analysis identifies extensive heterogeneity in the cellular composition of mouse Achilles tendons. Am. J. Physiol. Cell Physiol. 319, C885–C894

27. Gaut, L. and Duprez, D. (2016) Tendon development and diseases. Wiley Interdiscip. Rev. Dev. Biol. 5, 5–23

28. Calve, S., Dennis, R. G., Kosnik, P. E. 2nd, Baar, K., Grosh, K., and Arruda, E. M. (2004) Engineering of functional tendon. Tissue Eng. 10, 755–761

29. Calve, S., Lytle, I. F., Grosh, K., Brown, D. L., and Arruda, E. M. (2010) Implantation increases tensile strength and collagen content of self-assembled tendon constructs. J. Appl. Physiol. 108, 875–881

30. Novakova, S. S., Mahalingam, V. D., Florida, S. E., Mendias, C. L., Allen, A., Arruda, E. M., Bedi, A., and Larkin, L. M. (2018) Tissue-engineered tendon constructs for rotator cuff repair in sheep. J. Orthop. Res. 36, 289–299

31. de Wreede, R. and Ralphs, J. R. (2009) Deposition of collagenous matrices by tendon fibroblasts in vitro: a comparison of fibroblast behavior in pellet cultures and a novel three-dimensional long-term scaffoldless culture system. Tissue Eng. Part A 15, 2707– 2715

32. Schiele, N. R., Koppes, R. A., Chrisey, D. B., and Corr, D. T. (2013) Engineering cellular fibers for musculoskeletal soft tissues using directed self-assembly. Tissue Eng. Part A 19, 1223–1232

33. Mubyana, K. and Corr, D. T. (2018) Cyclic uniaxial tensile strain enhances the mechanical properties of engineered, scaffold-free tendon fibers. Tissue Eng. – Part A 24, 1808–1817

34. Percie du Sert, N., Ahluwalia, A., Alam, S., Avey, M. T., Baker, M., Browne, W. J., Clark, A., Cuthill, I. C., Dirnagl, U., Emerson, M., Garner, P., Holgate, S. T., Howells, D. W., Hurst, V., Karp, N. A., Lazic, S. E., Lidster, K., MacCallum, C. J., Macleod, M., Pearl, E. J., Petersen, O. H., Rawle, F., Reynolds, P., Rooney, K., Sena, E. S., Silberberg, S. D., Steckler, T., and Würbel, H. (2020) Reporting animal research: Explanation and elaboration for the ARRIVE guidelines 2.0. PLoS Biol. 18, e3000411

35. Park, N. R., Shetye, S. S., Bogush, I., Keene, D. R., Tufa, S., Hudson, D. M., Archer, M., Qin, L., Soslowsky, L. J., Dyment, N. A., and Joeng, K. S. (2021) Reticulocalbin 3 is involved in postnatal tendon development by regulating collagen fibrillogenesis and cellular maturation. Sci. Rep. 11, 10868

36. Eyre, D. (1987) Collagen cross-linking amino acids. Methods Enzymol. 144, 115–139

37. Fung, A., Sun, M., Soslowsky, L. J., and Birk, D. E. (2022) Targeted conditional collagen XII deletion alters tendon function. Matrix Biol. plus 16, 100123

38. Leiphart, R. J., Pham, H., Harvey, T., Komori, T., Kilts, T. M., Shetye, S. S., Weiss, S. N., Adams, S. M., Birk, D. E., Soslowsky, L. J., and Young, M. F. (2022) Coordinate roles for collagen VI and biglycan in regulating tendon collagen fibril structure and function. Matrix Biol. plus 13, 100099

39. Sun, M., Luo, E. Y., Adams, S. M., Adams, T., Ye, Y., Shetye, S. S., Soslowsky, L. J., and Birk, D. E. (2020) Collagen XI regulates the acquisition of collagen fibril structure, organization and functional properties in tendon. Matrix Biol. 94, 77–94

40. Kallenbach, J. G., Freeberg, M. A. T., Abplanalp, D., Alenchery, R. G., Ajalik, R. E., Muscat, S., Myers, J. A., Ashton, J. M., Loiselle, A., Buckley, M. R., van Wijnen, A. J., and Awad, H. A. (2022) Altered TGFB1 regulated pathways promote accelerated tendon healing in the superhealer MRL/MpJ mouse. Sci. Rep. 12, 3026

41. Farhat, Y. M., Al-Maliki, A. A., Easa, A., O’Keefe, R. J., Schwarz, E. M., and Awad, H. A. (2015) TGF-β1 Suppresses Plasmin and MMP Activity in Flexor Tendon Cells via PAI-1: Implications for Scarless Flexor Tendon Repair. J. Cell. Physiol. 230, 318–326

42. Farhat, Y. M., Al-Maliki, A. A., Chen, T., Juneja, S. C., Schwarz, E. M., O’Keefe, R. J., and Awad, H. A. (2012) Gene expression analysis of the pleiotropic effects of TGF-β1 in an in vitro model of flexor tendon healing. PLoS One 7, e51411

43. Tan, G.-K., Pryce, B. A., Stabio, A., Brigande, J. V, Wang, C., Xia, Z., Tufa, S. F., Keene, D. R., and Schweitzer, R. (2020) Tgfβ signaling is critical for maintenance of the tendon cell fate. Elife 9

44. Hou, Y., Mao, Z., Wei, X., Lin, L., Chen, L., Wang, H., Fu, X., Zhang, J., and Yu, C. (2009) The roles of TGF-beta1 gene transfer on collagen formation during Achilles tendon healing. Biochem. Biophys. Res. Commun. 383, 235–239

45. Pryce, B. A., Watson, S. S., Murchison, N. D., Staverosky, J. A., Dünker, N., and Schweitzer, R. (2009) Recruitment and maintenance of tendon progenitors by TGFbeta signaling are essential for tendon formation. Development 136, 1351–1361

46. Havis, E., Bonnin, M. A., De Lima, J. E., Charvet, B., Milet, C., and Duprez, D. (2016) TGFβ and FGF promote tendon progenitor fate and act downstream of muscle contraction to regulate tendon differentiation during chick limb development. Dev. 143, 3839–3851

47. Kaji, D. A., Howell, K. L., Balic, Z., Hubmacher, D., and Huang, A. H. (2020) Tgfβ signaling is required for tenocyte recruitment and functional neonatal tendon regeneration. Elife 9

48. Ansorge, H. L., Adams, S., Birk, D. E., and Soslowsky, L. J. (2011) Mechanical, compositional, and structural properties of the post-natal mouse Achilles tendon. Ann. Biomed. Eng. 39, 1904–1913

49. Li, X., Pongkitwitoon, S., Lu, H., Lee, C., Gelberman, R., and Thomopoulos, S. (2019) CTGF induces tenogenic differentiation and proliferation of adipose-derived stromal cells. J. Orthop. Res. Off. Publ. Orthop. Res. Soc. 37, 574–582

50. Liu, J., Tao, X., Chen, L., Han, W., Zhou, Y., and Tang, K. (2015) CTGF positively regulates BMP12 induced tenogenic differentiation of tendon stem cells and signaling. Cell. Physiol. Biochem. Int. J. Exp. Cell. Physiol. Biochem. Pharmacol. 35, 1831–1845

51. Linderman, S. W., Shen, H., Yoneda, S., Jayaram, R., Tanes, M. L., Sakiyama-Elbert, S. E., Xia, Y., Thomopoulos, S., and Gelberman, R. H. (2018) Effect of connective tissue growth factor delivered via porous sutures on the proliferative stage of intrasynovial tendon repair. J. Orthop. Res. Off. Publ. Orthop. Res. Soc. 36, 2052–2063

52. Desai, S. and Jayasuriya, C. T. (2020) Implementation of Endogenous and Exogenous Mesenchymal Progenitor Cells for Skeletal Tissue Regeneration and Repair. Bioeng. (Basel, Switzerland) 7

53. Szczesny, S. E. and Corr, D. T. (2023) Tendon cell and tissue culture: Perspectives and recommendations. J. Orthop. Res. Off. Publ. Orthop. Res. Soc.

